# Engineered neuron-targeting, placental mesenchymal stromal cell-derived extracellular vesicles for *in utero* treatment of myelomeningocele

**DOI:** 10.1101/2021.09.22.461362

**Authors:** Xinke Zhang, Hongyuan Chen, Kewa Gao, Siqi He, Zhao Ma, Ruiwu Liu, Dake Hao, Yan Wang, Priyadarsini Kumar, Lalithasri Ramasubramanian, Christopher D Pivetti, Yuanpei Li, Fuzheng Guo, Fengshan Wang, Randy Carney, Diana L Farmer, Aijun Wang

**Author notes:** Correspondence author Aijun Wang, PhD, Surgical Bioengineering Laboratory, Department of Surgery, School of Medicine, University of California, Davis, 4625 2nd Ave., Research II, Suite 3005, Sacramento, CA, 95817, USA. Contributed equally (Xinke Zhang, Hongyuan Chen).

## Abstract

This study investigated the feasibility and efficiency of neuron-targeting hybrid placental mesenchymal stromal cell-derived extracellular vesicles (PMSC-EVs), engineered by membrane fusion with Targeted Axonal Import (TAxI) peptide modified, TrkB agonist 7,8-DHF-loaded liposomes for treatment of myelomeningocele (MMC) via intra-amniotic cavity administration. The prepared TAxI modified liposomes with 7,8-DHF were used to fuse with PMSC-EVs. Different fusion approaches were investigated and freeze-thaw-extrude method was found to be the optimal. The engineered PMSC-EVs had a uniform particle size and efficiently loaded 7,8-DHF. It also had typical markers of native EVs. Freeze-thaw-extrude process did not change the release profile of 7,8-DHF from engineered EVs compared to TAxI modified, 7,8-DHF loaded liposomes. The engineered EVs could elicit TrkB phosphorylation depending on the incorporation of 7,8-DHF while native EVs did not. The engineered EVs increased neurite outgrowth of apoptotic cortical neurons induced by staurosporine, suggesting that they exhibited neuroprotective function. In a rodent model of MMC, neuron-targeting, engineered EVs became an active targeting delivery system to MMC defect sites. Pups treated with engineered EVs had the lowest density of apoptotic cells and displayed a therapeutic outcome. The study suggests the potential use of engineered hybrid, active neuron-targeting EVs for the *in utero* treatment of MMC.

## 1 Introduction

Myelomeningocele (MMC), the most severe form of spina bifida, is a congenital neural tube defect that occurs during early fetal development, leading to the incomplete closure of the spinal column.^[1]^ Children born with MMC live with lifelong paralysis, bowel and bladder dysfunction, musculoskeletal deformities, and severe cognitive disabilities.^[2]^ The current standard of care for MMC is *in utero* skin closure over the exposed spinal cord during the second trimester of pregnancy.^[3]^ Despite the promising results of fetal intervention in patients with MMC, which decreases the risk of hindbrain herniation and need for cerebrospinal fluid shunting, improvements in motor function are still needed.^[4]^

Our lab has previously shown that placental derived mesenchymal stromal cells (PMSCs) have distinct neuroprotective abilities in both fetal ovine and rodent models of MMC.^[3, 5]^ *In utero* PMSC transplantation induced significant distal motor function rescue in the fetal ovine MMC model and led to a significant decrease in the number of apoptotic neurons in the retinoic acid induced rat MMC model. Particularly, in our fetal ovine studies of *in utero* MMC repair with PMSCs, we confirmed that PMSC transplantation successfully preserved large neurons in the spinal cord but PMSCs did not persist after *in utero* transplantation.^[3]^ Therefore, it is likely that the main regenerative effects of PMSCs result from paracrine mechanisms mediated by secreted factors, including extracellular vesicles (EVs) such as such as exosomes, microvesicles, and other membrane-bound secreted vesicles.^[6]^ EVs, are now recognized to play an important role in cell-to-cell communication by transporting various functional molecules, including proteins, lipids, nucleic acids, and other small molecules,^[7]^ all of which play a prominent role in regulating the behaviors of the targeted cells. Our laboratory has previously characterized PMSC-EVs and evaluated their neuroprotective function.^[8]^ PMSC-EVs have excellent neuroprotective activity *in vitro* and impart their neuroprotective effect likely via galectin 1.^[8]^ PMSC-EVs contain numerous proteins and RNAs involved in neuronal survival and development, such as hepatocyte growth factor (HGF), galectin 1, α-catenin, β-cateninplatelet-derived growth factor (PDGF), and integrins by proteomic and RNAseq analyses. PMSCs secrete significantly higher levels of brain-derived neurotrophic factor (BDNF) and HGF, compared with adult bone marrow-derived MSCs (BM-MSCs), and we previously demonstrated that these neurotrophic factors likely play a significant role in mediating the distinct neuroprotective effect of PMSCs.^[3]^ BDNF, a cognate ligand for the tyrosine kinase receptor B (TrkB) receptor, mediates neuronal survival, differentiation, synaptic plasticity, and neurogenesis.^[9]^ Yet, from our previous proteomic analysis, we confirmed that PMSC-EVs do not contain BDNF.^[8]^ This lack of BDNF expression might lead to a decrease of the neuroprotective effect of PMSC-EVs. Therefore, incorporating BDNF into PMSC-EVs may represent an effective approach to retain the superior neuroprotective function of PMSCs. However, clinical trials using recombinant BDNF have not succeeded, presumably because of its poor pharmacokinetic profile, short half-life, and other limitations in delivery.^[9, 10]^ BDNF mimics that possess better biocompatibility and longer half-life could solve these issues. Recently 7,8-dihydroxyflavone(7,8-DHF) was identified as a bioactive high-affinity TrkB agonist that stimulates autophosphorylation, receptor dimerization, and downstream signaling cascades, imitates BDNF and displays a powerful therapeutic effect for the treatment of various neurological diseases.^[9, 11–14]^ Here we proposed to engineer PMSC-EVs to encapsulate 7,8-DHF for improved neuroprotective function for the treatment of MMC.

EVs have attracted increasing attention as a promising drug delivery platform ^[15, 16]^ for a variety of regenerative medicine applications, due to their distinct features such as good biocompatibility, relatively long circulating half-life, unique ability to cross the blood-brain barrier (BBB), and lack of inherent toxicity. However, their intrinsic targeting ability to specific cell types is limited.^[17]^ Modification of EV surface with effective targeting molecules represents an effective approach to optimize and tune the interaction and uptake of EVs by the recipient cells, thus improving targeting efficiency of EVs to neurons.^[18]^ Here we accomplished this by loading PMSC-EVs with 7,8-DHF and a neuron targeted recombinant bacteriophage peptide, targeted axonal import (TAxI), which can deliver protein cargo into spinal cord motor neurons.^[19]^ Importantly, TAxI-Cre recombinase fusion proteins could induce selective recombination and tdTomato-reporter expression in motor neurons. TAxI peptide has been shown to co-localize with motor neurons in human tissue, ^[19]^ indicating the potential that modification of EV surfaces with TAxI may improve EV’s neuron targeting potential.

In this study, we tested methods to incorporate 7,8-DHF into PMSC-EVs and successfully modified PMSC-EV surfaces with TAxI. Direct incubation or modification with other agents to EVs have been reported.^[15, 17, 20–24]^ Yet, considering the limited production of EVs and inability for wholly bottom-up synthesis of EVs, liposome-EV fusion is more feasible. ^[25]^ In this study, we thoroughly investigated the technical parameters involved in the EV-liposome membrane fusion approach. TAxI modified liposomes encapsulating 7,8-DHF were first prepared by film dispersion and then the engineered hybrid EVs were obtained by fusing EVs and liposomes using an optimized freeze-thaw-extrude method. By this novel membrane-engineering strategy, we successfully encapsulated 7,8-DHF into hybrid EVs and modified the surface of EVs with TAxI peptide (Figure 1). Additionally, modified and loaded hybrid EVs could provide sustained release of 7,8-DHF. The engineered PMSC-EVs displayed superior neuroprotective effects *in vitro* and *in vivo* compared to native EVs. We further validated the therapeutic targeting potential of the engineered, hybrid, neuron-targeting PMSC-EVs in a fetal rodent MMC model.

**Figure 1.**
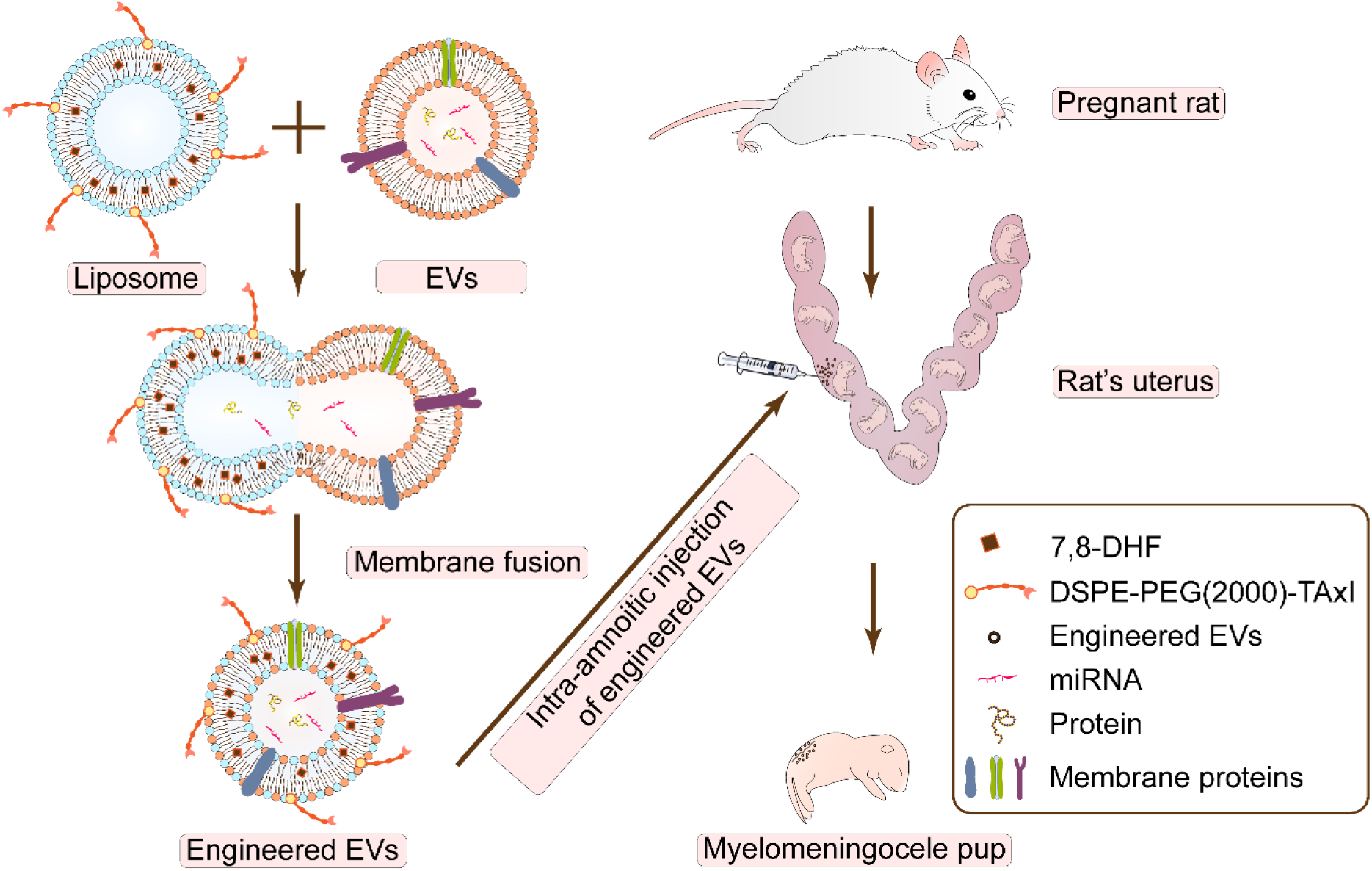
The schematic illustration of engineering EVs with 7,8-DHF and TAxI modification. Engineered EVs were generated by membrane fusion of TAxI modified, 7,8-DHF loaded liposomes with native PMSC-EVs. TAxI modification was designed to target exposed neurons in the MMC defect and loading 7,8-DHF was designed to provide neuroprotective effect. Biodistribution and therapeutic efficacy were further investigated in the retinoic acid-induced fetal rat MMC model via intra-amniotic injection of the engineered EVs.

## 2. Materials and methods

### 2.1 Culture of PMSCs and isolation of PMSC-EVs

We obtained PMSCs from our lab PMSC banks that have been well characterized for their multipotency and surface marker expression as previously described. ^[26]^ PMSCs at passage 4 were seeded in T175 flask at a density of 20,000 cells/cm^2^ and cultured for 48h in DMEM high glucose, 5% EV-depleted fetal bovine serum (FBS) (Thermo Fisher Scientific, Waltham, MA, USA), 20ng/ml EGF (AdventBio, Elk Grove Village, IL, USA), 20 ng/mL basic FGF, 100 U/mL penicillin and 100 μg/mL streptomycin at 37 °C, 5% CO2. Conditioned media was collected and EVs were isolated by differential centrifugation as described previously.^[8]^ Briefly, media was sequentially centrifuged at 300 *x g* for 10 min, 2000 *x g* for 20 min and passed through 0.2 μm filter. The media was concentrated and centrifuged at 8836 *x g* using the SW28 rotor and Beckman Coulter L7 ultracentrifuge. To collect EVs, the supernatant was centrifuged at 112,700 *x g* for 90 min, and the pellet was resuspended in phosphate buffered saline (PBS) and spun again.^[8]^ The final pellet was resuspended in 10 μL of PBS per flask used and stored at −80 °C.

### 2.2 Synthesis of DSPE-PEG-TAxI and preparation of TAxI modified, 7,8-DHF loaded liposomes

Cyclized TAxI peptide sequence (SACQSQSQMRCGGG) was used as described earlier.^[19]^ Reactive dibenzylcyclootyne (DBCO) group was linked at the amino group of glycine at the N-terminus of TAxI peptide via DBCO-NHS in presence of diisopropylethylamine. G-SACQSQSQMRCGGG was synthesized using standard Fmoc solid-phase peptide synthesis and cyclized with CLEAR OX resin as previously reported.^[27, 28]^ It was purified through *high-performance liquid chromatography (HPLC)* with a Waters auto-purification system (Milford, MA, USA) consisting of a 2998 detector, a 2525 pump, and a 2767 sample manager on a XTerra MS C18 OBD Prep Column (125Å, 5 µm, 19 × 150 mm) against an acetonitrile gradient. The identity was confirmed by matrix assisted laser desorption ionization-time of flight mass spectrometry (MALDI-TOF-MS) with a Bruker UltraFlextreme MALDI TOF/TOF (Billerica, MA, USA). Then DBCO-G-SACQSQSQMRCGGG was conjugated with DSPE-PEG2000 Azide (AVANTI POLAR LIPIDS, Inc) in acetonitrile by using biorthogonal copper-free Click chemistry. The reaction lasted for 3h at room temperature and was subsequently dialyzed and lyophilized. DSPE-PEG-G-TAxI was verified by MALDI-TOF-MS.

Thin film dispersion method^[29]^ was used to encapsulate 7,8-DHF into the TAxI modified liposomes. Briefly, 50.62μmol of soy phospholipid (AVANTI POLAR LIPIDS Inc., Alabaster, Alabama, USA), 7.76μmol of cholesterol (AVANTI POLAR LIPIDS Inc. Alabaster, Alabama, USA), 2.31μmol of DSPE-PEG-G-TAxI and 5.80μmol of 7,8-DHF (Millipore Sigma, Burlington, MA, USA) were mixed in 12mL solvent with chloroform and ethyl acetate (5:1 v/v). The solution was then evaporated under vacuum at 45°C for 2h to generate a thin film on the surface of round bottomed flask and subsequently hydrated by gentle mixing in 3mL of 10mM HEPES buffer overnight. Lipid extrusion was performed using an mini-extruder (AVANTI POLAR LIPIDS Inc., Alabaster, Alabama, USA) by passing the sample through filters with 0.4μm porosity. Liposomal purification was accomplished with Amicon Ultra-4 centrifugal filter units with a 100kDa MWCO (Millipore Sigma, Burlington, MA, USA) to eliminate free 7,8-DHF in the mixture. To calculate encapsulation efficiency, encapsulated 7,8-DHF in liposomal preparation was released by permeabilization of liposomes with methanol. The amount of 7,8-DHF (total and encapsulated) was determined spectrophotometrically at 300 nm using a standard curve ranging from 10μg/mL to 50μg/mL. The percent of encapsulation was calculated based on the ratio of encapsulated 7,8-DHF over the total 7,8-DHF used during the preparation.

### 2.3 Screening parameters to produce engineered EVs

#### 2.3.1 Fluorescent labeled liposome preparation

Liposomes were prepared and purified using the same method described in section 2.2 using fluorescently labeled lipid NBD-PE (AVANTI POLAR LIPIDS Inc., Alabaster, Alabama, USA) and Rhod-PE (AVANTI POLAR LIPIDS Inc., Alabaster, Alabama, USA) in the mixture. The molecules were dissolved in organic solvent chloroform: ethyl acetate (5:1 v/v) using the following molarities: 50.62μmol for soy phospholipid, 7.76μmol for cholesterol, 2.31μmol for DSPE-PEG-G-TAxI, 5.80μmol for 7,8-DHF, 0.065μmol NBD-PE and 0.065μmol Rhod-PE. The final concentration of fluorescent NBD-PE and Rhod-PE in liposomes was 0.0216mM, and the concentration of 7,8-DHF in liposomes was 1.9×10^3^μM.

#### 2.3.2 Screening the ratios of EVs and fluorescent liposomes

Fluorescent liposomes containing 1.9×10^3^μM 7,8-DHF, labeled with 0.0216mM NBD-PE and 0.0216mM Rhod-PE, described in previous section, were diluted 10-, 100- and 1000-fold in 10mM HEPES buffer. EVs (1×10^11^ EVs/mL) were then mixed with the diluted liposomes by 1:1 volume respectively. The membranes were fused using the freeze-thaw method, ^[25]^ where the mixture was flash frozen in liquid nitrogen and thawed at room temperature for 10min and repeating the freeze-thaw cycle 6 times. Fusion efficiency was evaluated using established FRET assays.^[25]^ Briefly, fusion was monitored using a fluorescence couple of NBD-PE and Rhod-PE within the lipid bilayer membrane. The fluorescence of the mixture was measured using a SpectraMax^®^ i3 Imaging Cytometer (Molecular Devices, San Jose, CA, USA) or an FP8000 fluorescence spectrometer (JASCO, Tokyo, Japan). Excitation of NBD-PE at 460 nm induces fluorescence emission at 534 and 592 nm, which corresponds to NBD-PE and Rhod-PE emissions, respectively. Lipid dilution due to membrane fusion increases fluorescence intensity at 534nm and decreases the intensity at 592 nm. FRET dissolution efficiency (E_FD_) of the hybrid EV-liposome mixture was defined as *E*_*FD*_ = *F*_534_/ (*F*_534+_ *F*_592_), where *F*_534_ and *F*_592_ represent the fluorescent intensities at 534 and 592 nm, respectively.

#### 2.3.3 Screening of membrane fusion parameters to generate hybrid EVs

Four hybridization methods were investigated. (1) EVs and liposomes were mixed and incubated in a water bath at 37°C for 2h. (2) EVs and liposomes were mixed, sonicated and incubated in a water bath at 37°C for 2h. (3) EVs and liposomes were mixed and fused by the freeze-thaw method. EVs and liposomes were mixed, frozen in liquid nitrogen and thawed at room temperature for 10min; this cycle was repeated 6 times. (4) EVs and liposomes were mixed and fused by the freeze-thaw-extrude method. After the EVs and liposomes were mixed and freeze-thawed 6 times, they were passed through filters with porosity 0.4μm on a mini-extruder. The fusion efficiency was evaluated using established FRET assays, as described above.

### 2.4 Characterization of engineered EVs

The surface markers of EVs and hybrid EVs were analyzed by Western blotting, as described previously.^[8]^ Briefly, 10μL of EVs or engineered EVs were treated with either NUPAGE LDS sample buffer (Thermo Fisher Scientific, Waltham, MA, USA) containing reducing agent dithiothreitol (DTT) for detecting Alg-2 interacting protein X (ALIX), Tumor Susceptibility gene 101 (TSG101), calnexin, or without DTT for detecting CD9 and CD63 proteins, and heated to 90°C.^[8]^ The samples were run, transferred, and probed with 1:500 dilution of primary antibodies (ALIX, CD9, TSG101 (Millipore Sigma, Burlington, MA, USA), calnexin (Cell Signaling Technology Inc., Danvers, MA, USA), CD63 (Thermo Fisher Scientific, Waltham, MA, USA), and imaged.

The size distributions and polydispersity index (PDI) of the engineered EVs were characterized with dynamic light scattering (DLS, Zeta sizer, Nano ZS) (Malvern Instruments Ltd., Worcestershire, UK.). Isolated PMSC-EVs were also visualized by nanoparticle tracking analysis (NTA) to detect EV size distribution and numbers using the Nano Sight LM10 (Malvern, Malvern, UK), which uses a 404 nm laser and sCMOS camera, as described previously.^[8]^

The morphology of engineered EVs was observed using an established negative staining protocol.^[30]^ Briefly, a 10μL drop of EVs was placed onto the grid in a dust-free space for 10 min. The excess was wicked off with filter paper and a 10μL drop of 2% uranyl acetate was applied to the grid, then was wicked off immediately. Grids air-dried thoroughly before viewing in a FEI Talos L120C electron microscope (FEI/Philips Inc, Hillsborough, OR, USA) at 80keV.

### 2.5 Accumulated drug release of hybrid engineered EVs

To determine the drug release profile of hybrid engineered EVs, TAxI modified 7,8-DHF liposomes and 7,8-DHF solution (control group) each containing 983μM of 7,8-DHF were prepared and loaded into dialysis cartridges (Millipore Sigma, Burlington, MA, USA). The cartridges were submerged into 1000 mL PBS (pH 7.4) stirred with moderate speed at ambient temperature. The 7,8-DHF in liposomes that retained in the dialysis cartridge was drawn with a microsyringe at multiple time points, and quantitatively measured by the UV absorbance of 7,8-DHF as described in section 2.2. The cumulative release, indicating the proportion of total 7,8-DHF being released from engineered EVs or TAxI modified 7,8-DHF liposomes was calculated and plotted over time.

### 2.6 Primary cortical neuron culture

Primary cortical neurons were collected from embryonic day 16 (E16) to E18 C57 mouse as described previously.^[31]^ The pregnant mouse was euthanized by CO2 asphyxiation and the neocortices of the embryos were collected in a HBSS buffer ice bath (Thermo Fisher Scientific, Waltham, MA, USA). The embryos were taken out and immersed in a dish containing 50mL of HBSS and 500μL of 15% glucose placed on ice. Embryonic brain cortices were collected, removed of their meninges, minced by trituration with 1mL pipette, and digested in papain (Worthington Biochemical Corporation, Lakewood, NJ, USA) containing DNase I (Millipore Sigma, Burlington, MA, USA) at 37°C for 1h. After the incubation, the suspension was passed through a 100μm filter and spun at 300 *x g* for 10min. The cells then underwent two sequential enzyme incubation and centrifugation steps. The final cell pellet was resuspended in neuron-plating medium containing DMEM, 10 % fetal bovine serum, 2 mM L-glutamine, 100 IU/mL penicillin and 0.10 mg/mL streptomycin, pH 7.4. The cells were seeded onto poly-L-lysine (Millipore Sigma, Burlington, MA, USA) coated plates at a cell density of 5× 10^5^ cells/cm^2^ and cultured at 37 °C, with 5 % CO_2_ and 95 % humidity. After 4 h of seeding, the medium was changed to Neurobasal medium (Thermo Fisher Scientific, Waltham, MA, USA) supplemented with B-27 (Thermo Fisher Scientific, Waltham, MA, USA), 0.2 mM L-glutamine (Thermo Fisher Scientific, Waltham, MA, USA), 100 IU/mL penicillin, and 0.10 mg/mL streptomycin. The culture medium was changed every two days. As a marker of cortical neurons, MAP2 expression was detected by immunofluorescent staining using anti-MAP2 antibodies (Cell Signaling Technology Inc., Danvers, MA, USA). The cells used for the following experiments were aged between 6 and 9 days of culture.

### 2.7 Engineered EVs elicited TrkB activation in cortical neurons

TAxI modified, 7,8-DHF loaded liposomes containing 250nM of 7,8-DHF, 1×10^8^ PMSC-EVs, and hybrid engineered EVs containing PMSC EV number of 1×10^8^ and 250nM 7,8-DHF were incubated with cortical neurons for 30min. Cortical neurons were fixed in 10% formalin for 10 min, and 1% bovine serum albumin (BSA) in 1X DPBS was used to block non-specific binding sites for one hour at RT. The cells were then stained with anti-phospho-TrkB Tyr515 (Thermo Fisher Scientific, 1:200) antibody in DPBS containing 1% BSA and incubated overnight at 4 °C. Cells were then incubated with Alexa594 conjugated secondary antibody (Thermo Fisher Scientific, 1:500) for 1 h at room temperature. The cells were counterstained with DAPI (1:5000) for 5 min and mounted with Prolong Diamond Antifade Mountant (Thermo Fisher Scientific, Waltham, MA, USA). A Nikon laser-scanning confocal microscope was used to acquire confocal images. The intensity of *p*-Trkb was calculated by Image J software.

### 2.8 Neuroprotection assay and caspase-3 acitvity for engineering EVs

Primary cortical neurons were seeded onto glass bottom 8-well μ-slides (ibidi GmbH, Matinsried, Germany). The neurons were pre-incubated with either 1×10^8^ PMSC-EVs, TAxI modified liposomes containing 250nM of 7,8-DHF or engineered EVs containing PMSC EV number of 1×10^8^ and 250nM 7,8-DHF for 30min. Cells were treated with 1μM staurosporine for 4 h to induce apoptosis. After apoptosis induction, nanoparticles in concentration described above, were added to cells and incubated for 20h at 37 °C, 5% CO2. After 20h the cells were washed twice with PBS. 2μM Calcein AM (Thermo Fisher Scientific, Waltham, MA, USA) in PBS was added to the cells and stained for 2 min. The cells were then imaged at 5 random positions per well at 5X magnification using a motorized Carl Zeiss Axio Observer D1 inverted microscope. To analyze neurite outgrowth and manually count live cells, the images were processed by WimNeuron Image Analysis (Wimasis, Cordoba, Spain). Additionally, caspase-3 activity was determined by Caspase-3 Activity Assay Kit (Cell Signaling Technology, Inc., Danvers, MA, USA). Primary cortical neurons were seeded onto 96-well plates. EVs, 7,8-DHF liposomes and engineered EVs, in the amount described previously, were added to primary neurons before staurosporine induction and incubated for 30 min. After staurosporine induction, EVs, 7,8-DHF liposomes and engineered EVs were added to cells again and incubated for 4h at 37 °C, 5% CO2. After 4h, the caspase-3 activity was determined.

### 2.9 Targeting MMC defects in the Retinoic acid-induced rat MMC model

To characterize the biodistribution *in utero*, liposomes and EVs were labeled with fluorescent DiR (Invitrogen™ Molecular Probes™, Thermo Fisher Scientific, Waltham, MA, USA). To label PMSC-EVs, native PMSC-EVs were mixed with DiR and were subjected to the freeze-thaw process to obtain a high incorporating efficiency. DiR-labeled TAxI modified liposomes were prepared with the addition of DiR into the mixture of lipid. The preparation method was the same as the non-fluorescent liposomes, described in section 2.2. DiR-labeled native PMSC-EVs and DiR-labeled liposomes served as the control groups. The concentration of DiR in liposomes and hybrid, engineered PMSC-EVs were both 22.5μg/mL.

The animal protocol was approved by the University of California Davis Institutional Animal Care and Use Committee (IACUC) and was performed according to the criteria provided by the Institute for Laboratory Animal Research Guide for Care and Use of Laboratory Animals. Via oral gavage, time mated Sprague–Dawley rats (Envigo, Livermore, CA) were dosed with 40 mg/kg of all-trans retinoic acid (RA) (Millipore Sigma, Burlington, MA, USA) on embryonic age (EA) 10 days (EA10) between a 2 hr.^[5]^ To initiate the biodistribution study, DiR-labeled nanoparticles were injected into the intra-amniotic cavity on EA20. 5% inhalational isoflurane was used to induce anesthesia, and was maintained at 1%–3% during surgery. Dams were placed supine on a temperature regulated warming pad set to 38°C. A midline laparotomy was performed and the bicornuate uterus was exposed. A 32-G needle on a 100-μL syringe (Hamilton Company, Reno, NV) was introduced into every amniotic cavity containing a viable fetus via the ventral aspect of the fetus, avoiding the fetus, placenta, and umbilical cord. Each injection consisted of 25μL either engineered hybrid EVs or native PMSC-EVs or TAxI modified liposomes. The uterus was then placed back in the abdomen and the incision was closed in two layers. Then, at 2h post injection, pups were collected via terminal cesarean section and taken images was conducted using In Vivo Imaging Spectrum (IVIS) system (PerkinElmer, Waltham, MA, USA) with an excitation band-pass filter at 748nm and an emission wavelength of 800nm. The fluorescence intensity was represented as radiant efficiency in the region of interest (ROI). The targeting efficiency (E_ts_)=radiant efficiency in the region of MMC defect/radiant efficiency in all the regions.

### 2.10 In vivo activity in the Retinoic acid-induced rat MMC model

The dams received injection of different formulations into the intra-amniotic cavity on EA17 days. The administration method was described in section 2.9. Each injection consisted of 10×10^8^ nanoparticles (native EVs, liposomes, or hybrid engineered EVs) in 25μL PBS. The groups included: 1) native PMSC-EVs, 2) TAxI modified 7,8-DHF loaded liposomes, and 3) hybrid engineered EVs. Fetuses that received 25μL PBS or 25μL liposomes loaded with 2.5μg 7,8-DHF served as control groups. After injection, the uterus was returned to the abdomen and the incision was closed in two layers. 0.05 mg/kg Buprenorphine was used as pain medication and was administered twice daily 48h after surgery. On EA21, pups were collected via terminal cesarean section. The uterus was wholly exteriorized and the total number of live pups and the presence of previous intrauterine fetal demise was noted. Each pup was examined for presence of MMC and/or anencephaly. Only fetuses with isolated MMC were used for further analysis. Collected pups were subjected to intracardiac perfusion with PBS and were fixed in 10% formalin.

### 2.11. Histological analysis

Whole pup specimens were fixed with 10% formalin for 24 h, dehydrated in 30% sucrose for 72 h, and embedded in O.C.T compound (Sakura Finetek, Torrance, CA USA). 10μm serial cross sections were taken through the lumbosacral portion of the pup starting from the approximate midportion of the MMC defect.^[5]^ Hematoxylin and Eosin (H&E) staining was performed on the serial cross sections of the lumbosacral portion for each collected pup. Each slide was analyzed for the presence of spinal cord tissue. Samples without adequate spinal cord tissues were excluded from further analysis. Images were captured at 20X magnification and merged using a Keyence BZ9000 microscope (Keyence Corporation of America, Itasca, IL, USA).^[5]^ The cross-sectional height and width of the lumbosacral spinal cord were measured using ImageJ. Spinal cord compression (defined as the width/height ratio) was calculated to assess the degree of cord deformity.^[5]^ A total of 3 sections from each fetus were stained and 5 fetuses of each group were analyzed.

### 2.12 TUNEL (terminal deoxynucleotidyl transferase-mediated UTP end-labeling) assay

TUNEL assay (In Situ Cell Death Detection Kit, TMR-red, Roche Biochemical Reagents, Waltham, MA) was performed according to the manufacturer's specifications to stain the apoptotic cells.^[5]^ Images were captured at 20X magnification and merged using a Keyence BZ9000 microscope. Density of apoptotic cells was determined using Image J software by dividing the number of apoptotic cells per section by the cross-sectional spinal cord area of the section. A total of 3 sections at the epicenter of the defect from each fetus were collected and stained and 5 fetuses from each group were analyzed.

### 2.13 Statistical Analysis

Data are reported as mean ± standard deviation (SD) and differences were considered significant if *p*<0.05. Mann Whitney tests were used to compare differences of spinal cord compression (width/height) or apoptotic cell density between normal and MMC control groups. One way ANOVA was used to compare other data both *in vitro* and *in vivo* between different treatment groups with the control group.

## 3. Results

### 3.1 Synthesis of DSPE-PEG-G-TAxI and preparation of TAxI Modified, 7,8-DHF loaded liposomes

DSPE-PEG-G-TAxI was prepared by copper-free DBCO-azide Click chemistry (Figure 2A). DSPE-PEG-G-TAxI was measured by MALDI-TOF-MS,confirming that it was synthesized successfully. The average molecular weight of DSPE-PEG(2000) Azide, DBCO-G-TAxI, DSPE-PEG(2000)-TAxI were 2816, 1741 and 4538 Da and demonstrated that DSPE-PEG-G-TAxI was synthesized successfully (Figure 2 B&C).

**Figure 2.**
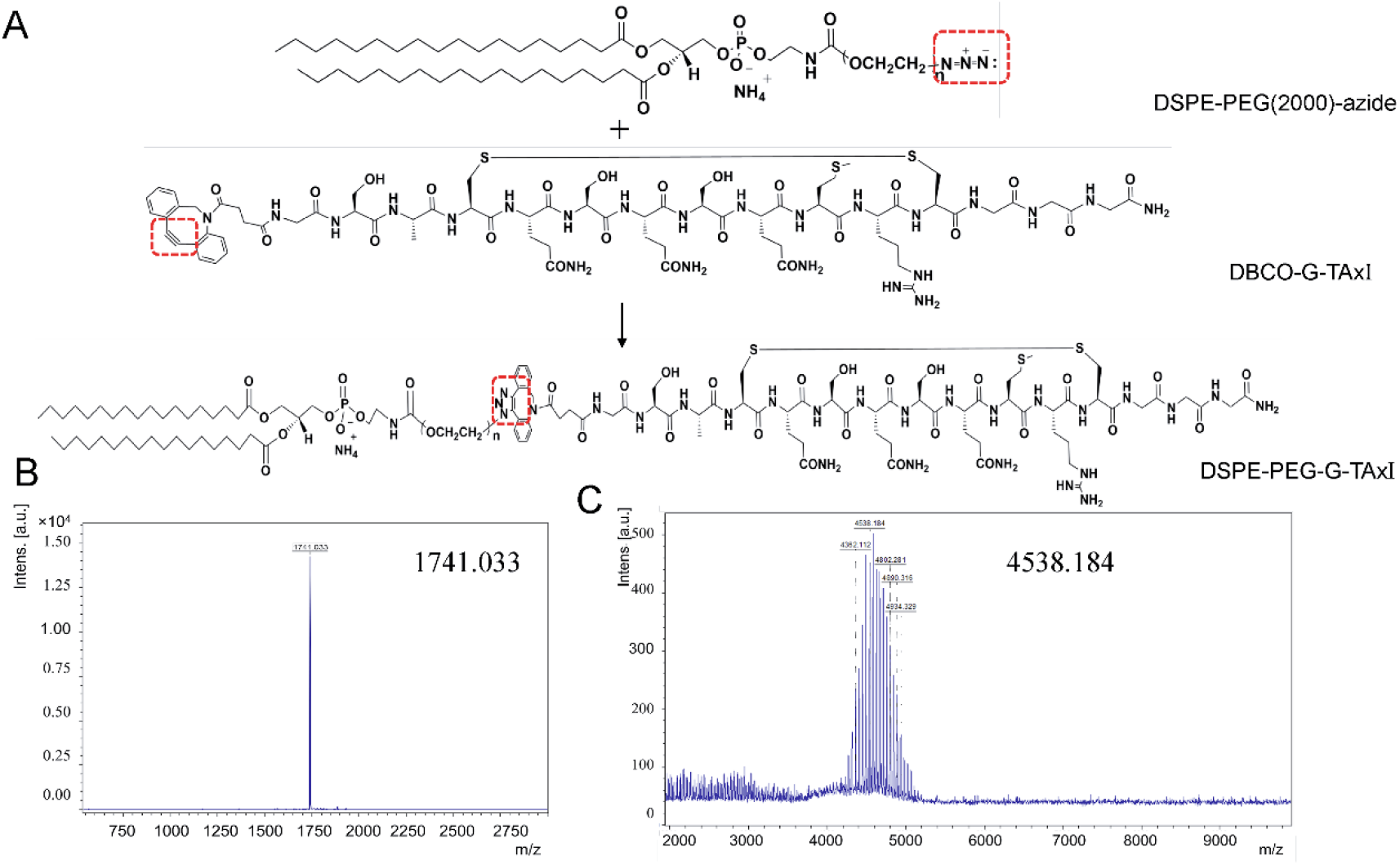
Synthesis and characterization of DSPE-PEG-G-TAxI. (A) Click chemistry reaction of DSPE-PEG2000 azide and DBCO-G-TAxI, (B) MALDI-TOF-MS result of DBCO-G-TAxI (mass = 1741.033), (C) MALDI-TOF-MS of DSPE-PEG-G-TAxI (Average mass = 4538.184)

DSPE-PEG-G-TAxI was used as one of the lipids in preparation of TAxI modified liposome loaded 7,8-DHF. The concentration of 7,8-DHF in original prepared liposomes was 1.9 mM and the entrapment efficiency was 89.2±3.9%, which was used in engineering hybrid EVs.

### 3.2 The screening of parameters to generate hybrid EVs with liposomes

First, we investigated the FRET dissolution efficiency at the hybrid ratio of EVs and fluorescent liposomes. The UV-Vis spectrum of mixture of EVs and fluorescent liposomes in different ratios are shown in Figure 3A-C. When the liposomes were diluted by 10 times, 100 times and 1000 times, E_FD_ of hybrid EVs were 0.11±0.019, 0.18±0.031, 0.217±0.043, respectively (Figure 3D) demonstrating an inverse relationship between the concentration of liposomes and E_FD_. To study the effect of different fusion methods on E_FD_, 10-fold, 100-fold and 1000-fold diluted 7,8-DHF, TAxI modified liposomes were used to hybrid EVs in parameter screening.

**Figure 3.**
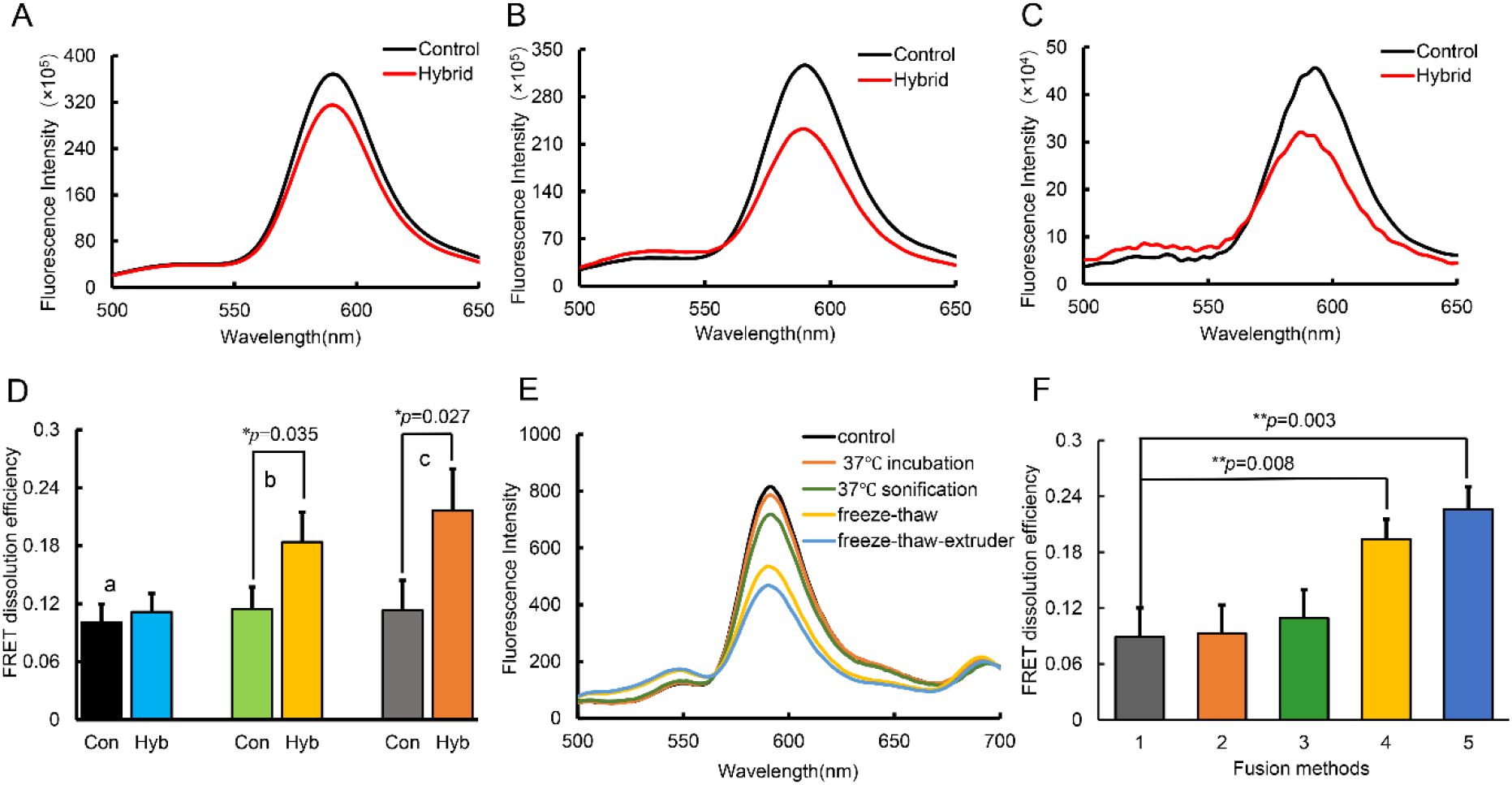
Screening parameters to produce hybrid engineered EVs by fusion of liposomes and PMSC-EVs and characterization of engineered EVs by FRET. (A-C) Fluorescence spectra of 10-fold,100-fold,1000-fold diluted fluorescent liposomes containing 1.9×10^3^μM 7,8-DHF fused with PMSC-EVs (1×10^11^EVs/ml) with excitation at 460 nm were characterized. The final concentration of 7,8-DHF liposomes used to engineer EVs was 1.9×10^2^μM (A), 19μM (B), 1.9μM (C), respectively. (D) FRET dissolution efficiency of mixture of 10-fold (a), 100-fold (b), 1000-fold (c) diluted liposomes with PMSC-EVs (Hyb), mixture with PBS (not EVs) served as control (Con). (E) Fluorescence spectra of fluorescent liposomes fused with PMSC-EVs by different methods. (F) FRET dissolution efficiency of fluorescent liposomes fused with PMSC-EVs by different methods-1(37 °C incubation), 2(37 °C sonification), 3(freeze-thaw),4(freeze-thaw-extrude) and 5 represents control. \**p*<0.05, \*\**p*<0.01, n = 3 repeated 3 times.

The UV-Vis spectrum of the mixture of EVs and fluorescent liposomes by different fusion methods are shown in Figure 3E-F. The FRET dissolution efficiency *E*_*FD*_ = *F*_534_/ (*F*_534_+ *F*_592_) in control group, 37°C incubation group, 37°C incubation-sonification group, freeze-thaw group and freeze-thaw-extrude group were 0.089±0.031, 0.093±0.0304, 0.1093±0.0302, 0.194±0.0212 and 0.226±0.0238, respectively. After the freeze-thaw-extrude process, *E*_*FD*_ was the highest. Since *E*_*FD*_ was the highest when using the freeze-thaw-extrude process, this approach was selected to conduct further experiments.

### 3.3 Characterization of engineered EVs

EVs were characterized according to the suggested experimental guidelines put forth by the International Society for Extracellular Vesicles.^[32]^ Western blotting analysis confirmed the presence of typical EV markers CD63, CD9, TSG101 and ALIX, and the absence of endoplasmic reticulum marker calnexin in engineered EVs (Figure 4A), which was the same as the native PMSC-EVs. PMSCs lysate was a cell control((Figure 4B). Size distribution showed that engineered EVs and native PMSC-EVs had a size range of 104.7±10.86 nm and 94.5±11.2 nm (Figure 4C). The 7,8-DHF entrapment efficiency of engineered EVs was 86.2±4.5%. Compared to liposomes, there was no obvious change of size and entrapment efficiency after freeze-thaw-extrude process. TEM images of the engineered EVs were shown in Figure 4D.

**Figure 4.**
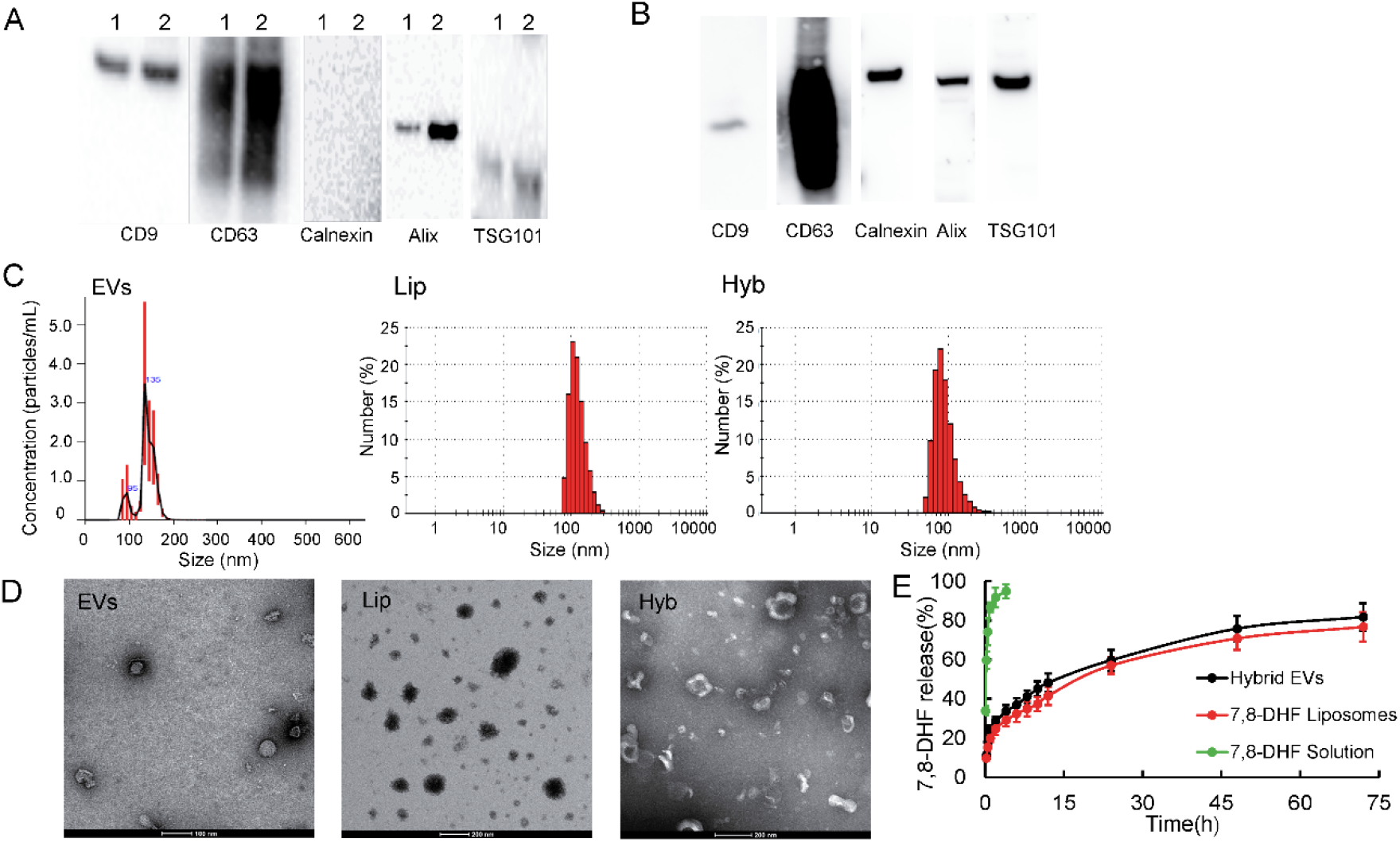
Characterization of engineered EVs and release profile of 7,8-DHF from liposomes and engineered EVs. (A) Western blot analysis of the expression of EV markers in the hybrid engineered EVs (lane 1) and native EVs (lane 2). (B) Western blot analysis of the expression of EV markers in the lysate of corresponding PMSCs as the cell control. (C) The size distribution of native PMSC-EVs (EVs), TAxI modified liposomes (Lip) and 7,8-DHF loaded hybrid engineered EVs (Hyb) by NTA. (D) TEM imaging of different nanoparticles. (E) The accumulative release profile of 7,8-DHF from 7,8-DHF solution (green line), TAxI modified, 7,8-DHF loaded liposomes (black line) and hybrid engineered EVs (red line).

### 3.4 Accumulated drug release of hybrid engineered EVs

Engineered hybrid EVs and TAxI liposomes could slowly release 7,8-DHF over time compared to 7,8-DHF solution. 7,8-DHF solution released 91.7±4.7% 7,8-DHF at 2h, while the engineered EVs released 24.6±3.2%, 56.9±4.2% and 76.7±7.5% at 2h, 24h, 72h respectively (Figure 4E). After the freeze-thaw-extrude process, the release profile of 7,8-DHF from engineered EVs did not change significantly.

### 3.5 The neuroprotection effect on primary cortical neurons

Over 95% of the isolated cortical neurons were positive for neuronal marker MAP2 (Support Information Figure S2). Both liposomes encapsulating 7,8-DHF and the 7,8-DHF loaded hybrid engineered EVs elicited TrkB phosphorylation shown by immunocytochemistry staining (Figure 5A) of phosphorylated TrkB (p-Trkb). Native PMSC-EVs did not elicit TrkB activation, indicating that the incorporated 7,8-DHF into liposomes or hybrid EVs was able to activate TrkB phosphorylation (Figure 5B), mimicking the function of BDNF and mediating neuroprotective activity on cortical neurons.

**Figure 5.**
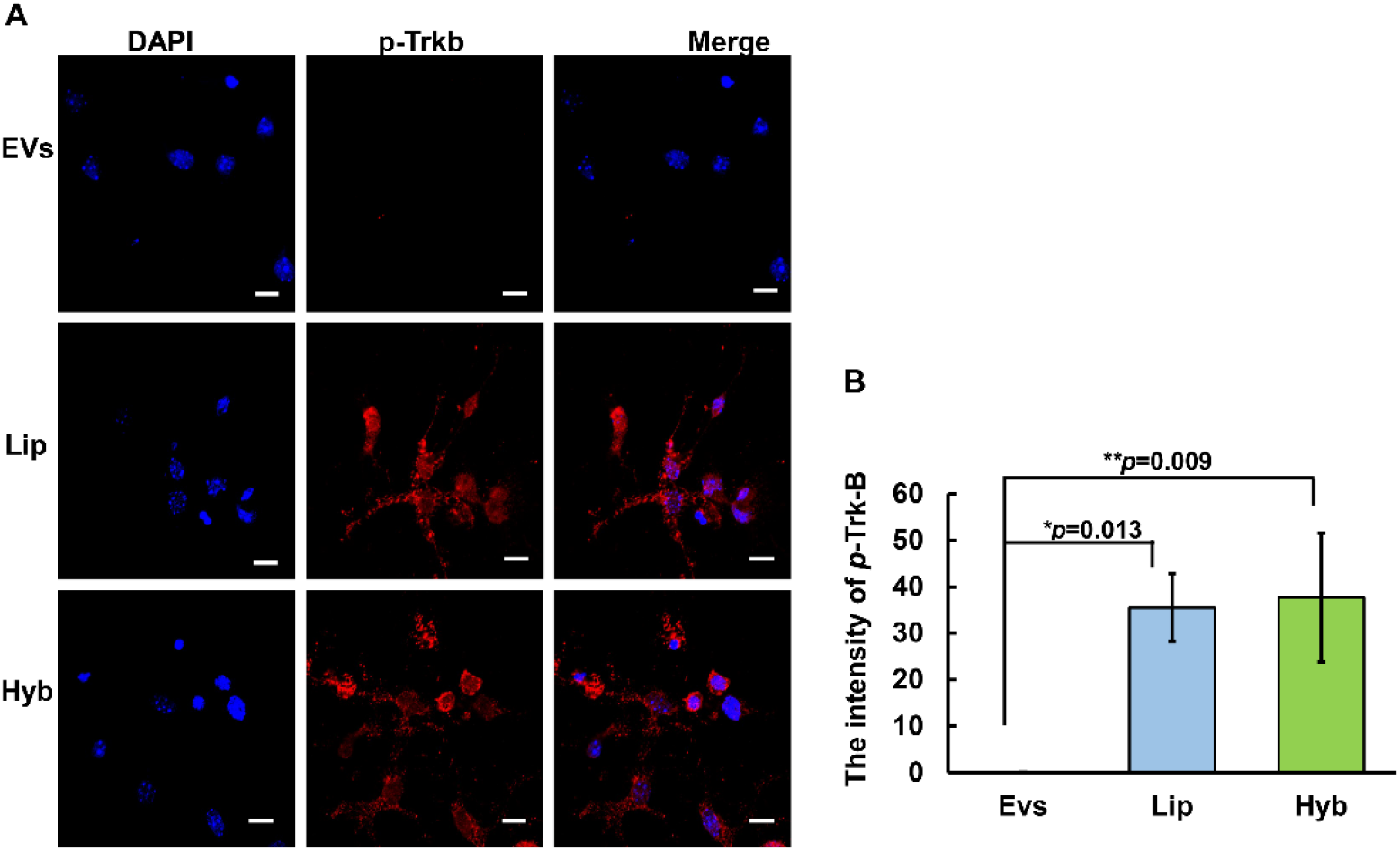
TrkB phosphorylation in primary cortical neurons by native EVs, 7,8-DHF loaded liposomes and hybrid engineered EVs. (A) Immunocytochemical staining of primary cortical neurons after treatment with native EVs (top panels), TAxI modified, 7,8-DHF loaded liposomes (middle panels) and the hybrid engineered EVs (lower panels). Representative images of cells stained with DAPI (blue, left panels), and phosphorylated TrkB (red, middle panels) and merged images (right panels). Both TAxI modified, 7,8-DHF loaded liposomes and the hybrid engineered EVs increased phosphorylation of TrkB compared to native EVs. Scale bar = 10μm. (B) Quantification of the fluorescent intensity of p-TrkB (red) in different groups. EVs, Lip, Hyb represent native PMSC-EVs, TAxI modified, 7,8-DHF loaded liposomes and hybrid engineered EVs, respectively. *\*p*<0.05, *\*\*p<*0.01, n=3

To study the neuroprotective function of engineered hybrid EVs, engineered hybrid EVs were added directly to primary cortical neuronal cells before and after the induction of apoptosis. Calcein AM imaging (Figure 6A) and WimNeuron image analysis (Figure 6B) showed that all groups, including PMSC-EVs (EVs), 7,8-DHF loaded liposomes (Lip), and hybrid engineered EVs (Hyb) displayed increased cortical neuronal cells, total branch points and longer circuitry and tube length compared to controls thus imparting a neuroprotective effect on apoptotic neurons (Figure 6C). Besides, treating primary neurons with EVs, liposomes and engineered EVs attenuated the increase in caspase-3 activity induced by staurosporine(Figure 6D), which further verified the neuroprotection function of 7,8-DHF liposomes, EVs and engineered EVs. Consistent with the WimNeuron image analysis results, there were no significant differences between nanoparticle treatment groups.

**Figure 6.**
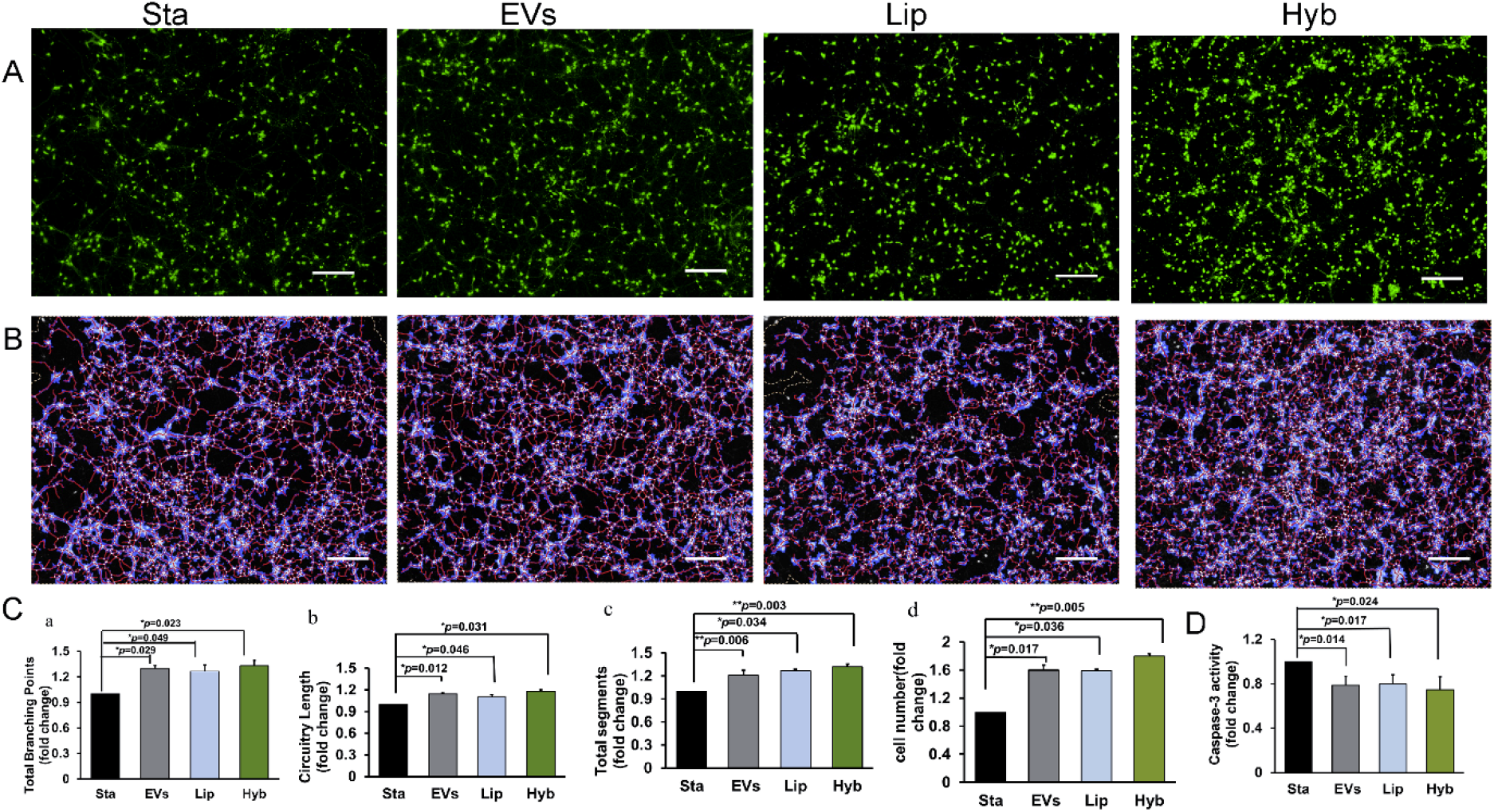
Neuroprotective effects of different nanoparticles on apoptotic primary cortical neurons. Staurosporine treatment was applied to primary cortical neurons were subjected to induce apoptosis. Before and after this treatment, primary neurons were incubated with different nanoparticles. (A) Calcein staining pictures of the cortical neurons subject to different treatments. (B) WimNeuron image analysis images of the neurons subject to different treatments. (C) WimNeuron image analysis results were quantified and demonstrated a significant increase in total branching points (a), circuitry length (b), total segments (c) and cell number (d) in the presence of different nanoparticles compared to the control (staurosporine treatment only). (D) Caspase-3 activity showed a significant suppression in the presence of EVs, liposomes and engineered EVs compared with the staurosporine group. Sta, EVs, Lip, Hyb represent staurosporine treatment group, native PMSC-EVs, TAxI modified, 7,8-DHF loaded liposomes and hybrid engineered EVs, respectively. \**p*<0.05, ** *p*<0.01. n=3 repeated 3 times. Scale bar = 200μm.

### 3.6 Specimen collection

28 dams were treated with 40 mg/kg of retinoic acid resulting in 380 total pups with a fetal viability rate of 66.05% at the time of collection on EA21 by Cesarean delivery (C-section). Of the 129 viable pups, 104 (41.43%) fetuses were normal, 105 (41.83%) fetuses presented with MMC defects, 39 (15.54%) had anencephaly, and 3 (0.12%) had gastroschisis. Of the pups that had MMC defect, the number of pups that were administered with PBS, hybrid engineered EVs, native PMSC-EVs, TAxI modified 7,8-DHF liposomes injection, DiR-labeled hybrid engineered EVs, DiR-labeled native EVs and DiR-labeled liposomes were 8, 10, 11, 14, 8, 7 and 9, respectively. 5 normal pups (no MMC) were served as normal control and 5 PBS injected MMC pups served as negative control. 5 MMC pups in each group were analyzed in analysis of the deformity of MMC spinal cords and density of apoptotic cells in the spinal cord.

### 3.7 Targeting to MMC defect

To evaluate the *in vivo* biodistribution of engineered EVs in fetal MMC rats, DiR-labeled hybrid engineered EVs, native PMSC-EVs, and TAxI modified, 7,8-DHF loaded liposomes were administered by intra-amniotic injection and the fetuses were imaged at 2h post injection using IVIS. The results indicated that the hybrid engineered EVs and TAxI modified liposomes had high targeting efficiency to the MMC defect at 54.0±20.6% and 59.2±19.6% respectively at 2h after administration compared to native PMSC-EVs with the targeting efficiency of 22.5±13.4% (Figure 7). Though the hybrid engineered EV group and native PMSC-EV group were not significantly different (*p*=0.089), the increased targeting to the MMC defect of engineered EVs after TAxI modification suggest that this modification may be beneficial for the treatment of MMC defect.

**Figure 7.**
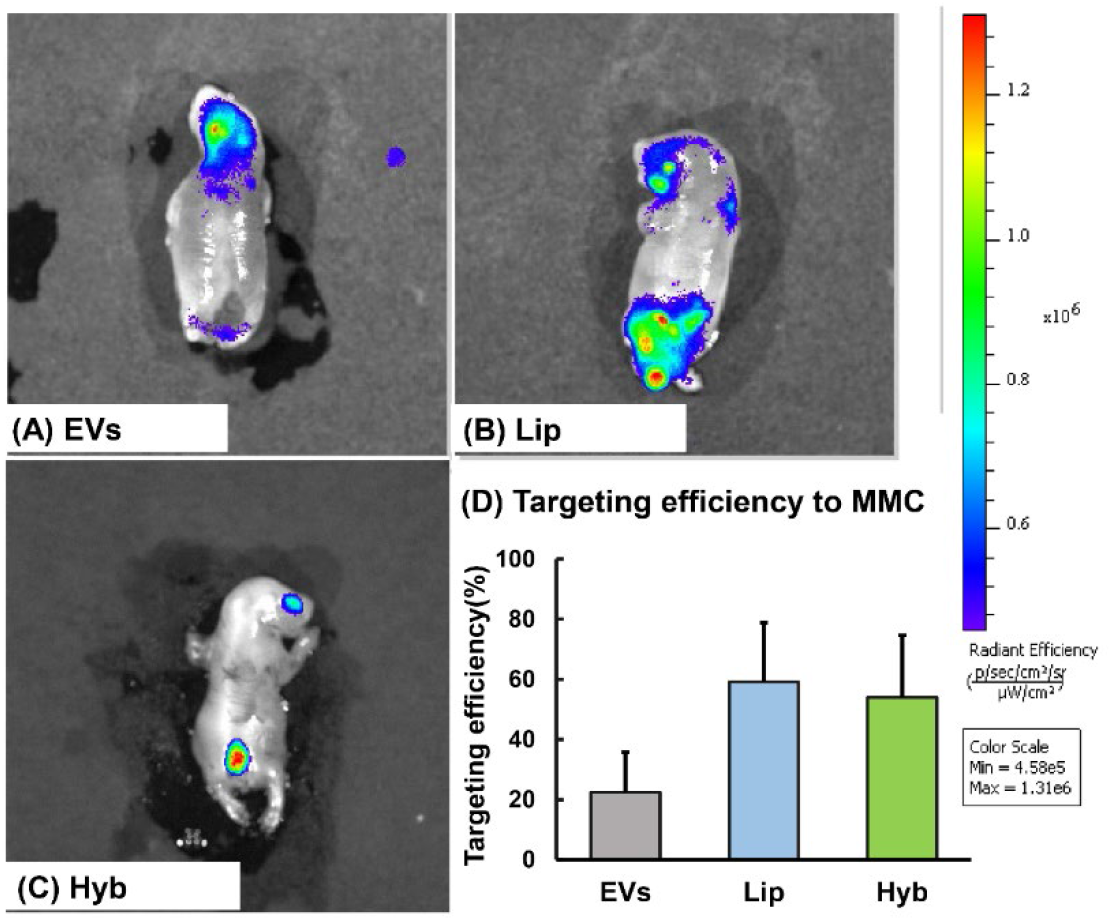
Targeting efficiency of different nanoparticles to the MMC defect. IVIS image of the MMC fetal rats collected by Cesarean delivery (C-section) on EA21 after treatment with DiR labeled native PMSC-EVs (A, EVs), DiR labeled TAxI modified 7,8-DHF loaded liposomes (B, Lip), and DiR labeled hybrid engineered EVs (C, Hyb). The targeting efficiency results (D) showed that though no statistical differences were identified due to large individual variations between individuals, TAxI modified 7,8-DHF loaded liposomes (Lip) and the hybrid engineered EVs (Hyb) showed a trend resulting in a higher targeting efficiency to the MMC defect compared to the native PMSC-EVs (EVs). EVs, Lip, Hyb represent fetal MMC groups treated with native PMSC-EVs, TAxI modified, 7,8-DHF loaded liposomes, and hybrid engineered EVs, respectively. n=3.

### 3.8 Analysis of the deformity of MMC spinal cords

The ratio of the cross-sectional width/height of the spinal cord was used as a parameter for spinal cord deformation and compression.^[5]^ A larger width/height ratio indicated greater spinal cord compression and deformation in the specimen.^[5]^ The mean width/height ratio of control MMC pups (5.921 ± 0.519) was significantly greater than the width/height ratio in normal pups (1.103 ± 0.036) (*p*<0.001) (Support Information Figure S5), suggesting a greater degree of spinal cord compression and deformation in MMC pups. The mean width/height ratio of all treatment groups were not significantly different compared to the control MMC group, suggesting that the treatments had no impact on cord compression.

### 3.9. Density of apoptotic cells in the spinal cord

To determine the density of apoptotic cells/mm^2^ of spinal cord tissue, TUNEL analysis was performed on representative samples from each study group. MMC control pups (PBS injection) showed significantly greater density of apoptotic cells (cells/mm^2^) (203.2±53.4), compared to normal pups (12.3 ±4.3) (*p*=0.008) (Figure 8). Pups treated with hybrid engineered EVs had the lowest density of apoptotic cells (111 ± 55.6), compared to other treatment groups and was significant compared to the MMC only control group (*p*=0.035). Pups treated with TAxI modified liposomes loaded with 7,8-DHF (124.4±43.5) also had significantly lower density of apoptotic cells compared to MMC only control group (*p*=0.02) (Figure 8). Native PMSC-EV treatment (136.6±37.7) also decreased the density of apoptotic cells compared to control group (*p*=0.044). Pups treated with engineered EVs had a slightly lower density of apoptotic cells than that of native PMSC-EVs, though there was no significant difference between them.

**Figure 8.**
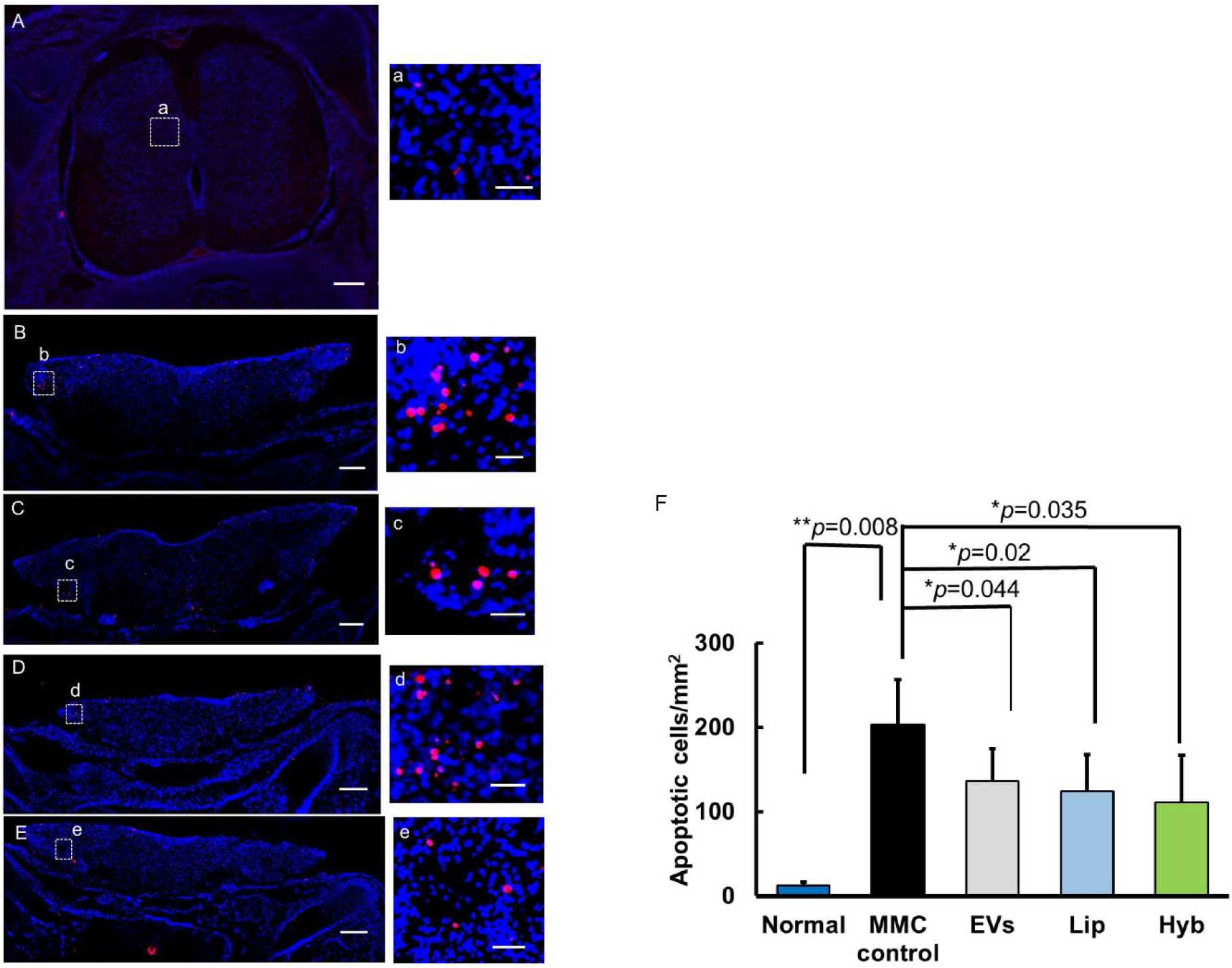
TUNEL analysis to evaluate density of apoptotic cells in the spinal cord tissues in fetal MMC rats. Red TUNEL positive cells are shown with blue DAPI nuclear counterstain. Representative images for spina cord of pups in normal group (A), untreated MMC group (B), EV treated MMC group (C), TAxI modified, 7,8-DHF loaded liposome treated MMC group (D), and hybrid engineered EV treated MMC group (E), (a, b, c, d, e) are the magnified areas in different groups. Statistical analysis of TUNEL positive cells (F) showed untreated MMC spinal cords had a significantly greater density of apoptotic cells (cells/mm^2^) compared to normal, untreated fetal rat spinal cords. MMC pups treated with PMSC-EVs, TAxI modified, 7,8-DHF loaded liposomes and the hybrid engineered EVs showed significant decrease in density of apoptotic cells compared to untreated MMC pups, n=5. *\*p*<0.05. Scale bar=200μm for (A, B, C, D, E), scale bar = 50μm for (a, b, c, d, e).

## 4. Discussion

There is a well-accepted “two-hit hypothesis” to explain MMC pathogenesis.^[1]^ The “first hit” describes the failure of neural tube closure, which is thought to lead to the spinal cord’s exposure to the intrauterine environment. The “second hit” describes the chemical and mechanical trauma sustained by the spinal cord due to its exposure to the intrauterine environment during pregnancy. This “second hit” results in the irreversible neurological impairments seen after birth. Interventions such as fetal surgical repair of the defect *in utero* protects the exposed spinal cord and prevents the “second hit” to a certain degree. However, in previous clinical trials it was shown that 58% of children remain unable to ambulate. *In utero* stem cell therapy has the potential to augment the current standard of care of *in utero* surgery and further prevent the second hit.^[4, 33]^ Recently, several investigators demonstrated the therapeutic potential of mesenchymal stromal cells (MSCs) derived from amniotic fluid, bone marrow, and embryonic stem cells (ESCs) using experimental MMC models.^[4, 33, 34]^ Our laboratory also confirmed that using PMSCs *in utero* transplantation can result in a significant improvement in neurologic function at birth in the fetal rodent and ovine models of MMC.^[3, 35–37]^ We confirmed that PMSCs did not persist after *in utero* transplantation.^[3]^ Therefore, it is thought that the main regenerative effects of PMSCs arise from paracrine secretion mechanisms, including EVs such as exosomes and microvesicles.^[6]^ EVs are now recognized to play an important role in intercellular communication by transporting various functional molecules, including proteins, lipids, microRNAs, and mRNAs^[7]^, all of which play a prominent role in regulating the behaviors of the targeted cells. Compared to stem cells, EVs have their own merits as acellular therapy, which avoids the risks of stem cell transplantation including low engraftment rate, immune response and tumor formation. Additionally, EVs are more convenient for storage and transportation. Though PMSC-EVs were validated to have neuroprotective effects *in vitro*, the therapeutic effect on MMC *in vivo* remains to be determined. In the present study, the activity of PMSC-EVs on MMC *in vivo* was investigated. To overcome some drawbacks such as lacking BDNF and targeting efficiency to neurons, engineered hybrid EVs were designed. These novel EVs were modified by TAxI, which could induce better targeting of EVs to neurons, and also encapsulated with 7,8-DHF, a BDNF mimic, potentially empowering EVs with better therapeutic neuroprotective effects. However, identifying the optimal administration dosage of EVs and 7,8-DHF represents the critical first step in translating this novel treatment for *in vivo* applications. Thus, in our preliminary experiments, in order to screen their optimal concentrations, the EVs or 7,8-DHF liposomes at different concentrations were injected into the intra-amniotic cavity of MMC rats respectively. 2×10^8^, 10×10^8^ EVs or 7,8-DHF solution equal to 2.5μg, 5μg, 7.5μg 7,8-DHF per fetus were studied. The results showed that treatment with 2×10^8^ EVs did not significantly reduce the density of apoptotic neurons compared to the control group. Pups treated with 10×10^8^ EVs had significantly lower density of apoptotic cells compared to the pups treated with MMC control. Pups with 7,8-DHF at dosage of 2.5μg, 5μg, 7.5μg had significantly lower density of apoptotic cells compared to the control group. But 7,8-DHF at dosage of 7.5μg had a higher demise rate (75.6%). In order to guarantee the therapeutic efficacy as well as safety, 10×10^8^ EVs and 7,8-DHF at dosage of 2.5μg per fetus in hybrid engineered EVs were selected to investigate *in vivo* experiments.

Due to their ideal characteristics and native structure, EVs are nanocarriers with great promise for clinical application. EVs can be engineered at the cellular level under natural conditions, but EV modification by exogenous strategies is also established.^[20]^ Compared with the cellular level endogenous methods, exogenous methods after production in cell culture expand the possible sources for EVs and might greatly facilitate mass production by mixing with other particles. Successful modification of EVs after isolation is informed by their formation mechanisms and structural characteristics. As EVs and liposomes are both composed of lipid membrane bilayers, they can be fused under certain conditions. From this, TAxI modified, 7,8-DHF loaded liposomes were first prepared to modify the surface of EVs and encapsulate 7,8-DHF into EVs. 7,8-DHF, a flavone derivative with high hydrophobicity, was incorporated into the lipid membrane by the thin film dispersing method with high encapsulating efficiency into liposomes. Limited water solubility of 7,8-DHF restricts its clinical use, which could be overcome by liposomal formulation. TAxI, a peptide which was demonstrated to deliver protein cargos into spinal cord motor neurons after intramuscular injection via a retrograde axonal transport mechanism, was used in this study as an active targeting peptide to deliver liposomes or EVs to neurons. DSPE-PEG-G-TAxI, synthesized with copper-free Click chemistry, was used to modify the surface of liposomes or exosomes as one lipid composition in order to realize the targeting efficiency to neurons. The biodistribution assay showed that TAxI modified liposomes or engineered EVs both displayed higher targeting efficiency to the MMC defect.

With respect to the engineered strategies, we investigated several methods such as incubating liposomes with EVs, sonicating the mixture, as well as freeze-thaw and extrude the mixture. These approaches have been used in producing engineered EVs, but little is known about the difference between them. In our research, the fused EV-liposomes were characterized using a FRET assay, and the freeze/thaw approach has been found as the most efficient method. However, after 6 freeze/thaw cycles the fused EV-liposome constructs aggregated greatly and some particles became very large and not uniform (Support information Figure S1). Thus, the engineered EVs were then extruded to obtain homogeneous nanoparticles. Furthermore, the ratio of EV-liposome was found to affect the fusion efficiency as well. As the ratio of EVs increased in mixture, fusion efficiency was much higher, indicating more exosomal lipid was incorporated into the liposomes. The engineered EVs by the freeze-thaw-extrude approach had a uniform size at (104.7±10.86) nm, high entrapment efficiency of 86.2±4.5% and maintained the EV markers similar to native EVs. Besides, freeze-thaw-extrude process did not change the release profile of 7,8-DHF from engineered EVs obviously and did not negatively impact the neuroprotective effect of PMSC- EVs *in vitro*. *In vitro* neuroprotective experiments, EVs, 7,8-DHF loaded liposomes, and engineered EVs all displayed neuroprotective effects on apoptotic neurons, compared to the control (staurosporine treatment). Yet, there was no statistical significance between the engineered EVs, liposomes and native PMSCs EVs. In the *in vitro* neuroprotection assay, 7,8-DHF liposomes, EVs and engineered EVs were directly added to the cultured neurons and they could all directly interact with the apoptotic neurons, and impart neuroprotective effects on the apoptotic neurons. Therefore, the advantages of engineered EVs could not possibly be fully displayed by *in vitro* experiments.

We further investigated the biodistribution and therapeutic effects of engineered PMSC-EVs on fetal MMC using a rat model. In the biodistribution experiments, engineered EVs, liposomes and native PMSC-EVs were all labeled with near-infrared dye DiR, which was incorporated into the lipophilic bilayer. Engineered EVs and TAxI modified liposomes both exhibited higher targeting efficiency to MMC defects after 2h intra-amniotic injection. We investigated the biodistribution at several time points (4h, 6h), but found weak fluorescence signal at 4h and 6h. This might be caused by quick metabolization of EVs or the concentration of DiR was too low to be detected at 4h or 6h. A larger width/height ratio of the spinal cord at the MMC defect was associated with greater spinal cord compression and specimen deformity. Intra-amniotic injection of nanoparticles did not change the spinal cord width/height ratio greatly. Compared to surgery repair, intra-amniotic injection treatment did not directly close the MMC defect, thus this indirect treatment cannot decrease width/height greatly. However, engineered EVs exhibited higher therapeutic efficiency via active targeting to exposed neurons (Figure 7). Engineered EVs resulted in the lowest density of apoptotic cells in the spinal cord tissue compared to other treatment groups. Furthermore, PMSC-EVs and 7,8-DHF liposomes also displayed neuroprotective activity in the rodent MMC model, which was the first investigation of their *in vivo* effects in an MMC model.

Yet, there was no significant difference between the hybrid EVs and native PMSC-EVs either in MMC targeting efficiency or in TUNEL analysis experiments. This was possibly caused by many factors. First, the complicated intrauterine environment and the status of developing fetuses have made it difficult to screen effective concentrations of EVs or 7,8-DHF. Though we preliminarily screened several concentrations of EVs and 7,8-DHF, the range of concentrations may not have been wide enough, therefore the chosen concentrations of EVs and 7,8-DHF could be further optimized. Secondly, the modification rate of TAxI could impact the targeting efficiency of hybrid EVs. Engineered EVs with higher modification percentage of TAxI could potentially deliver more nanoparticles to the MMC defect. The modification rate of TAxI on EVs and liposomes may need further optimization to impart effective targeting efficiency of the nanoparticles to the MMC defect. Thirdly, the fetal induction and treatment of MMC in pregnant rats are technically challenging. Due to the technical difficulties with the model, we have only tested limited numbers of MMC pups in this study, which had restricted us from evaluating the parameters extensively. Though engineered EVs did not show significant superiority in targeting and TUNEL analysis compared to native PMSC-EVs, the trend was obvious. In future studies, we plan to further optimize the concentrations of EVs and 7,8-DHF, as well as the modification rate of TAxI on engineered EVs to further improve the therapeutic efficacy of this innovative treatment.

With this novel EV engineering strategy and modification, we have demonstrated several enhancements to PMSC-EVs. First, the number of nanoparticles undoubtedly increased following fusion with liposomes, thereby boosting the productivity of artificial EVs. Secondly, the therapeutic efficiency was improved with incorporation of 7,8-DHF, and the poor water solubility issue of this potent agent was solved. Thirdly, with the incorporation of TAxI, a motor neuron specific targeting molecule, the engineered EVs have gained an active targeting ability towards motor neurons. All of these advantages make the innovative hybrid engineered EVs a more promising therapeutic modality for MMC.

However, there are other limitations to this research. The mechanism under which how TAxI exactly targets EVs to MMC defect via intra-amniotic injection is unclear. The accurate ratio of EVs and liposomes has not been studied thoroughly. The number of EVs or the total protein mass value in EVs is usually used to determine the concentration EVs. In our research, the number of EVs was used to determine the exosomal concentration for both the *in vitro* and *in vivo* experiments. Protein concentration in EVs was not determined. Additionally, given the limited availability of antibodies to analyze fetal rat neuronal tissue, histological analysis of the fetal rodent spinal cord was restricted. The most important aspect is that RA-induced MMC rats die immediately after birth, and thus, the long-term outcomes with respect to motor dysfunction cannot be studied using this model. Therefore, we aim to answer these questions in future studies, such as a more expansive and reliable histological analysis of the fetal rat spinal cords, and translation into an ovine MMC model to rigorously test and validate motor function data after EV treatment.

## 5. Conclusion

In this study, we have provided a novel, feasible and effective strategy to engineer EVs with exogenous agents and active targeting peptide via membrane fusion with liposomes. The engineered EVs were confirmed to effectively target the MMC defect and reduce the number of damaged neurons significantly in the rodent MMC model. Therefore, the approach could be used to obtain artificial EVs to efficiently load hydrophobic drug with high dissolution efficiency and our engineered EVs could be a novel treatment strategy for fetal MMC.

## Acknowledgements

This work was partially supported by the National Institutes of Health research grants 5R01NS100761-02 and 1R01NS115860-01A1; the Shriners Hospitals for Children Postdoctoral Fellowship, No. 84705-NCA-19; and the Shriners Hospitals for Children Research Grants, No. 85108-NCA-19 and 85135-NCA-21. The authors would like to thank the Campus Mass Spectrometry Facilities (supported by NIH S10 award S10OD018913) at UC Davis for obtaining the mass spectral data.

