## Supplementary material for "Engineered neuron-targeting, placental mesenchymal stromal cell-derived extracellular vesicles for *in utero* treatment of myelomeningocele": supplemtal file

**Support information**

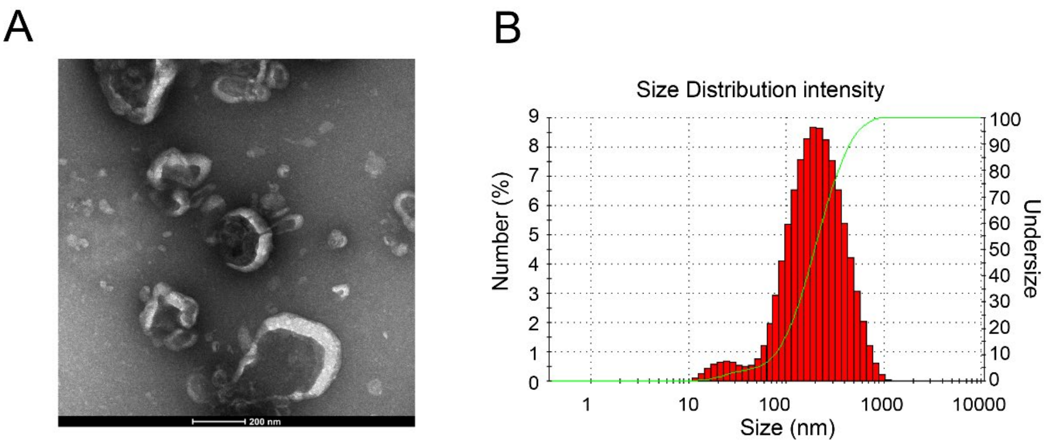


**Figure S1 TEM and size distribution characterization of engineered EVs subjected to freeze-thaw procedures**

After 6 freeze/thaw cycles the hybrid fused EV-liposome constructs aggregated greatly shown by TEM imaging (A) and the size of the particles became larger and less uniform characterized by NTA (B).

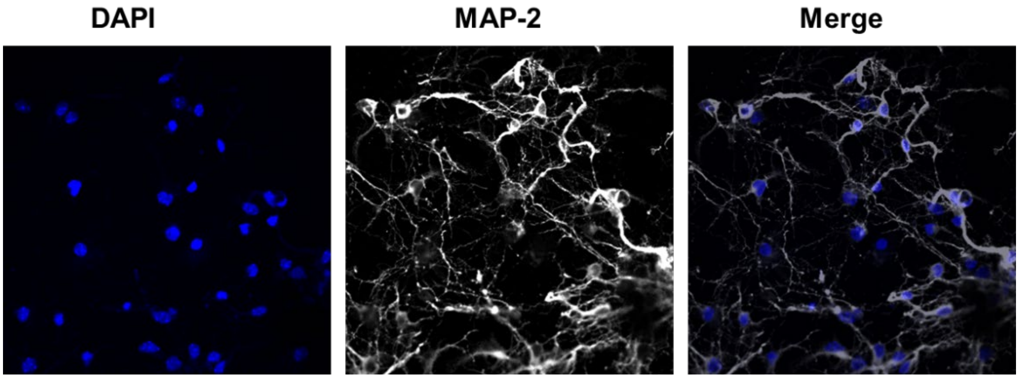


**Figure S2 Characterization of the isolated cortical neurons with immunofluorescent staining of neuronal marker MAP2.** Over 95% neurons express Map2 with DAPI nuclear counterstaining.

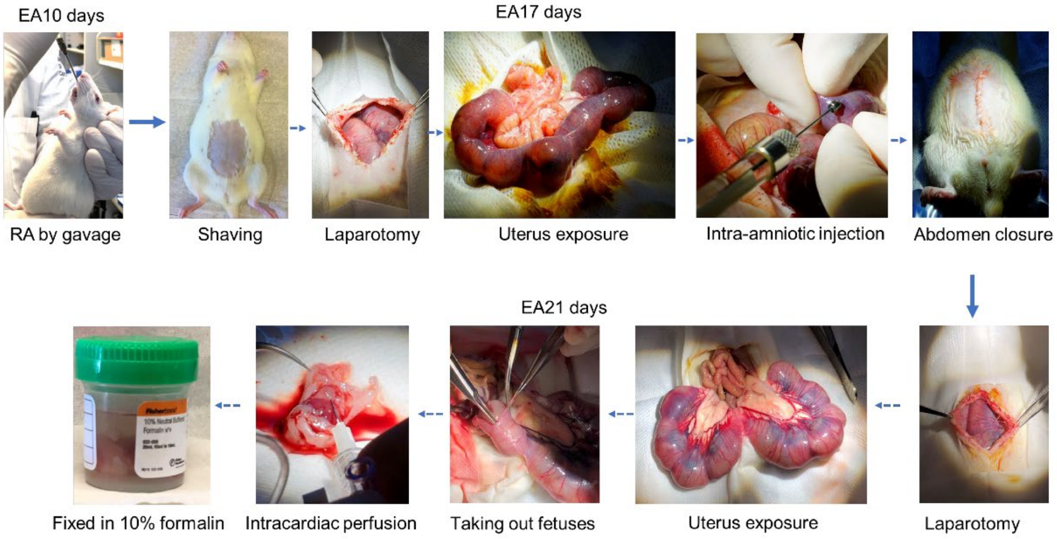


**Figure S3 The scheme of establishing the fetal rat MMC model, RA administration method and collection of the specimens.**

On EA10 day, RA was gavaged to the pregnant rat. On EA17 day, different preparations were intra-amniotic injected. On EA21 day, pups were collected after Cesarean delivery (C-section) followed by intracardiac perfusion.

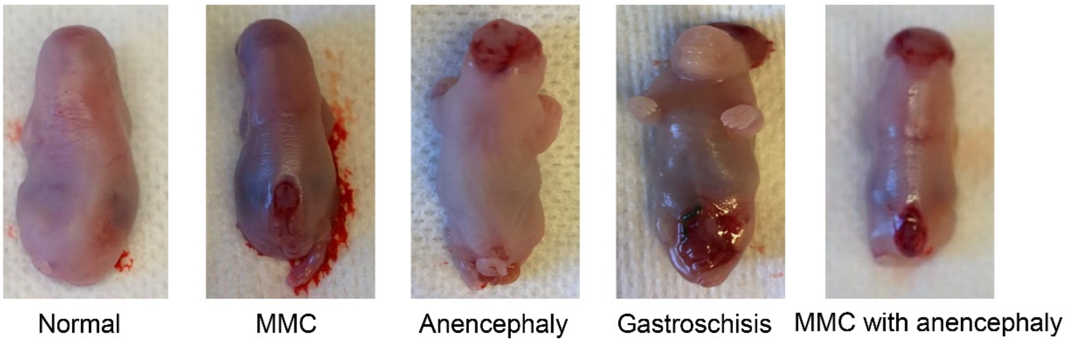


**Figure S4 Representative images of RA-induced abnormalities in fetal rats.**

Fetal rats were identified with abnormalities including myelomeningocele (MMC, spina bifida aperta), anencephaly, gastroschisis, MMC with anencephaly. Normal fetal rats was also shown for comparison.

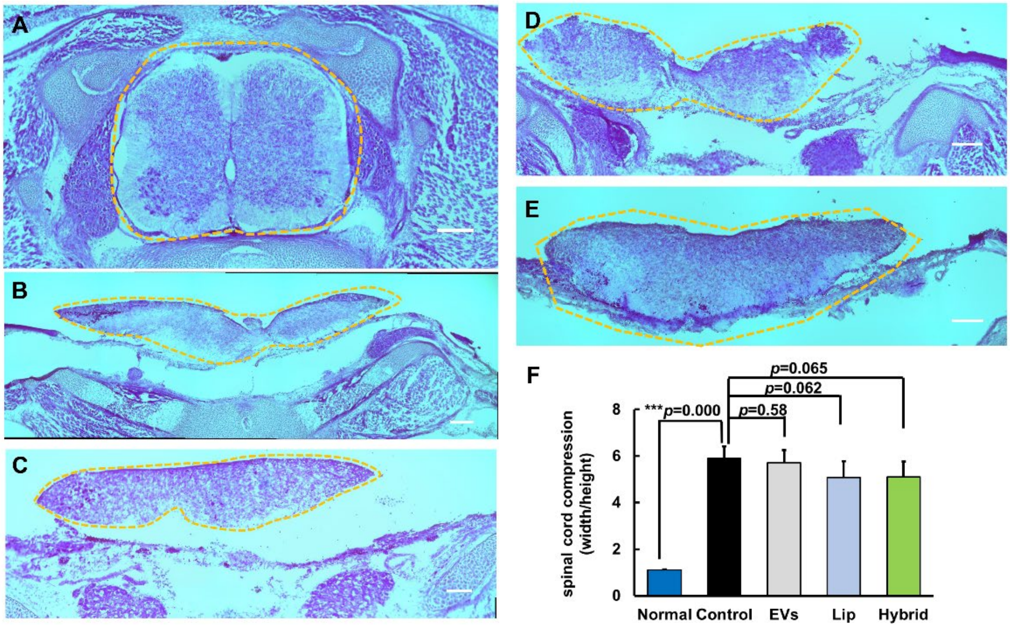


**Figure S5 Representative images of fetal rat spinal cord with HE staining**

(A) Normal, (B) untreated MMC, (C) native PMSC-EV treated MMC, (D) TAxI liposome treated MMC, (E) Hybrid engineered EV treated MMC. Spinal cord compression (width/height) results (F) showed that MMC pups with different treatments showed no significant difference compared to untreated MMC pups.
